# Overexpression of *ZmSTOP1-A* Enhances Aluminum Tolerance in Arabidopsis by Stimulating Organic Acid Secretion and Reactive Oxygen Species Scavenging

**DOI:** 10.1101/2023.09.05.556357

**Authors:** Chan Liu, Xiaoqi Hu, Lei Zang, Xiaofeng Liu, Yuhui Wei, Xue Wang, Xinwu Jin, Chengfeng Du, Yan Yu, Wenzhu He, Suzhi Zhang

**Affiliations:** Key Laboratory of Biology and Genetic Improvement of Maize in Southwest China of Agricultural Department, Ministry of Agriculture, Maize Research Institute, Sichuan Agricultural University, Chengdu 611130, China; Crop Research Institute, Sichuan Academy of Agricultural Sciences, Chengdu 610066, China

**Keywords:** *Zea mays*, Aluminum toxicity, *STOP1*, low pH, ZmSTOP1-A

## Abstract

Aluminum (Al) toxicity and low pH are major factors limiting plant growth in acidic soils. Sensitive to Proton Rhizotoxicity 1 (STOP1) transcription factor respond to these stresses by regulating the expression of multiple Al- or low pH-responsive genes. ZmSTOP1-A, a STOP1-like protein from maize (*Zea mays*), was localized to the nucleus and had transactivation activity. *ZmSTOP1-A* was expressed moderately in both roots and shoots of maize seedlings, but was not induced by Al stress or low pH. Overexpression of *ZmSTOP1-A* in Arabidopsis *Atstop1* mutant partially restored Al tolerance and completely low pH tolerance with respect to root growth. Regarding Al tolerance, *ZmSTOP1-A/Atstop1* plants showed clear upregulation of organic acid transporter genes, and leading to increased organic acid secretion and reduced Al accumulation in roots. Besides, the antioxidant enzyme activity in roots and shoots of *ZmSTOP1-A/Atstop1* plants was significantly enhanced, ultimately alleviating Al toxicity via scavenging reactive oxygen species. Similarly, ZmSTOP1-A could directly activate *ZmMATE1* expression in maize, positively correlated with the number of Al-responsive GGNVS *cis*-element in the *ZmMATE1* promoter. Our results revealed that ZmSTOP1-A antagonizes Al toxicity by enhancing organic acid secretion and reactive oxygen species scavenging in Arabidopsis, demonstrating that it is an important transcription factor conferring Al tolerance. Our findings help to comprehensively elucidate the role of STOP1-like transcript factor in enabling plants to detoxifying Al.

## Introduction

Acid soil accounting for 50% of potential arable land worldwide, yet can severely inhibit plant growth and reduce crop yields (Uexküll and Mutert, 1995). The primary limiting factor in acid soil is aluminum (Al) toxicity (Kochian et al., 2004). Below pH 5.0, Al solubilizes into the phytotoxic Al^3+^ form, which damages plant root structure, impair water and nutrient uptake, and ultimately inhibits normal plant growth (Kobayashi and Nishizawa, 2012; Kochian et al., 2015). Elucidating the genes involved in Al tolerance mechanisms is therefore crucial for improving crop yields through genetic approaches.

Generally, external exclusion and internal tolerance are two main mechanisms for plants to cope with Al stress (Ma et al., 2001). The external exclusion refers to the secretion of organic acid from the rhizosphere, of which chelate Al^3+^ and prevent its entry into the root. MATE (Multidrug and toxic compound extrusion) and AlMT1 (Aluminum activated malate transporter 1), two kinds of organic acid transporter of the external exclusion mechanism, are crucial for Al tolerance in many crops (Sasaki et al., 2006; Magalhaes et al., 2007). Internal tolerance involves sequestration of excessive Al into the vacuoles or immobilization in the cell wall. Additionally, recent evidence indicates scavenging of reactive oxygen species (ROS) also plays an important role in Al detoxification (Ranjan et al., 2021). Some genes including *ZmAT6* (Aluminum transporter 6), *AtGST* (Glutathione S-transferase) and *AtPrx64* (Peroxidases 64) have been implicated in removing ROS during Al stress (Ezaki et al., 2004; Wu et al., 2017; Du et al., 2020). To date, many factors including transcription factors, organic acid transporter, ABC transporter, Mg transporter, enzymes related to cell wall components or ROS scavenging, phytohormones, and small peptide have been reported to play roles in the Al stress response across plant species (Wei et al., 2021). Transcription factors, especially the STOP1-like class, are key regulators which charge the expression of many Al-responsive genes and play a central role in Al detoxification (Sawaki et al., 2009).

AtSTOP1, a C2H2 zinc finger transcription factor first indentified in Arabidopsis (Iuchi et al., 2007), has since been found in several crops including rice (*Oryza sativa)*(Yamaji et al., 2009), rice bean (*Vigna umbellata*) (Fan et al., 2015), and barley (*Hordeum vulgare*) (Wu et al., 2020). AtSTOP1 is a key regulator preventing Al toxicity by controlling the expression of 24 Al-responsive genes, including *AtALMT1* and *AtALS3* (Aluminum sensitive 3) (Sawaki et al., 2009). Its rice homolog OsART1 (Al resistance transcription factor 1) similarly targets 31 genes including *OsFRDL4* and *OsSTAR2* (Sensitive to aluminum rhizotoxicity 2) for Al detoxification. Notably, most of these AtSTOP1/ OsART1-regulated genes contain the *cis*-element GGN(T/g/a/C)V(C/A/g)S(C/G) in their promoters (Yamaji et al., 2009; Tsutsui et al., 2011). Beyond Al stress, STOP1-like transcription factor also respond to low pH by regulating a distinct set of genes, such as *AtSTOP2*, *AtGDH1* (Glutamate dehydrogenase 1), *AtCIPK23* (CBL-Interacting protein kinase 23) and *AtNRT1.1* (Nitrate transporter 1.1) (Sawaki et al., 2009; Ohyama et al., 2013; Ye et al., 2021). However, some STOP1 homologs are insensitive to low pH (Yamaji et al., 2009; Garcia-Oliveira et al., 2013; Che et al., 2018).

Maize is one of the most important food crops, yet its yield drops dramatically when grown on acidic soils (Lidon and Barreiro, 2002). The genetic mechanisms underlying Al toxicity response of maize remain poorly understood. To date, only six genes have been reported contributed to Al tolerance in maize, by facilitating citric acid efflux (*ZmMATE1* and *ZmMATE6*) (Maron et al., 2010; Du et al., 2021), ROS elimination (*ZmALDH1* and *ZmAT6*) (Du et al., 2020; Du et al., 2022), cell wall fixation of Al (*ZmXTH1*) (Du et al., 2021), and auxin transport (*ZmPGP1*) (Zhang et al., 2018). In this study, *ZmSTOP1-A* was first identified in maize and its Al-responsive expression pattern was explored. Most importantly, the roles of *ZmSTOP1-A* in Al- and low pH-tolerance were genetically investigated through ectopic expressing in the Arabidopsis mutant *Atstop1*. The mechanisms underlying *ZmSTOP1-A*-mediated Al tolerance was also examined by surveying expression of Al-responsive genes and measuring key physical indexes and Al content in the tested plants. This study provides important insights into how the maize *ZmSTOP1-A* responded to both Al toxicity and low pH. The findings expanded our understanding of the role STOP1-like transcription factors play in enabling crop species to adapt to acidic environments.

## Results

### Identification and phylogenetic analysis of ZmSTOP1-A

Like other STOP1-like transcription factors, the six ZmSTOP1s in maize contain four highly conserved putative C2H2 zinc finger domains characteristic of STOP1-like proteins (**Figure S1**). Phylogenetic analysis showed that ZmSTOP1-A and ZmSTOP1-B clustered on the same branch, closely related to SbSTOP1-d, three TaSTOP1s and AtSTOP1. In a branch above ZmSTOP1-A, ZmSTOP1-C and ZmSTOP1-D were most closely related to OsART1 and SbSTOP1-a, respectively. Meanwhile, ZmSTOP1-E and ZmSTOP1-F were in the same branch as AtSTOP2 and OsART2, adjacent to SbSTOP1-b and SbSTOP1-c (**Figure 1**). This reveals that STOP1-like proteins have complex evolutionary relationships characterized by both conservation and divergence.

**Fig. 1.**
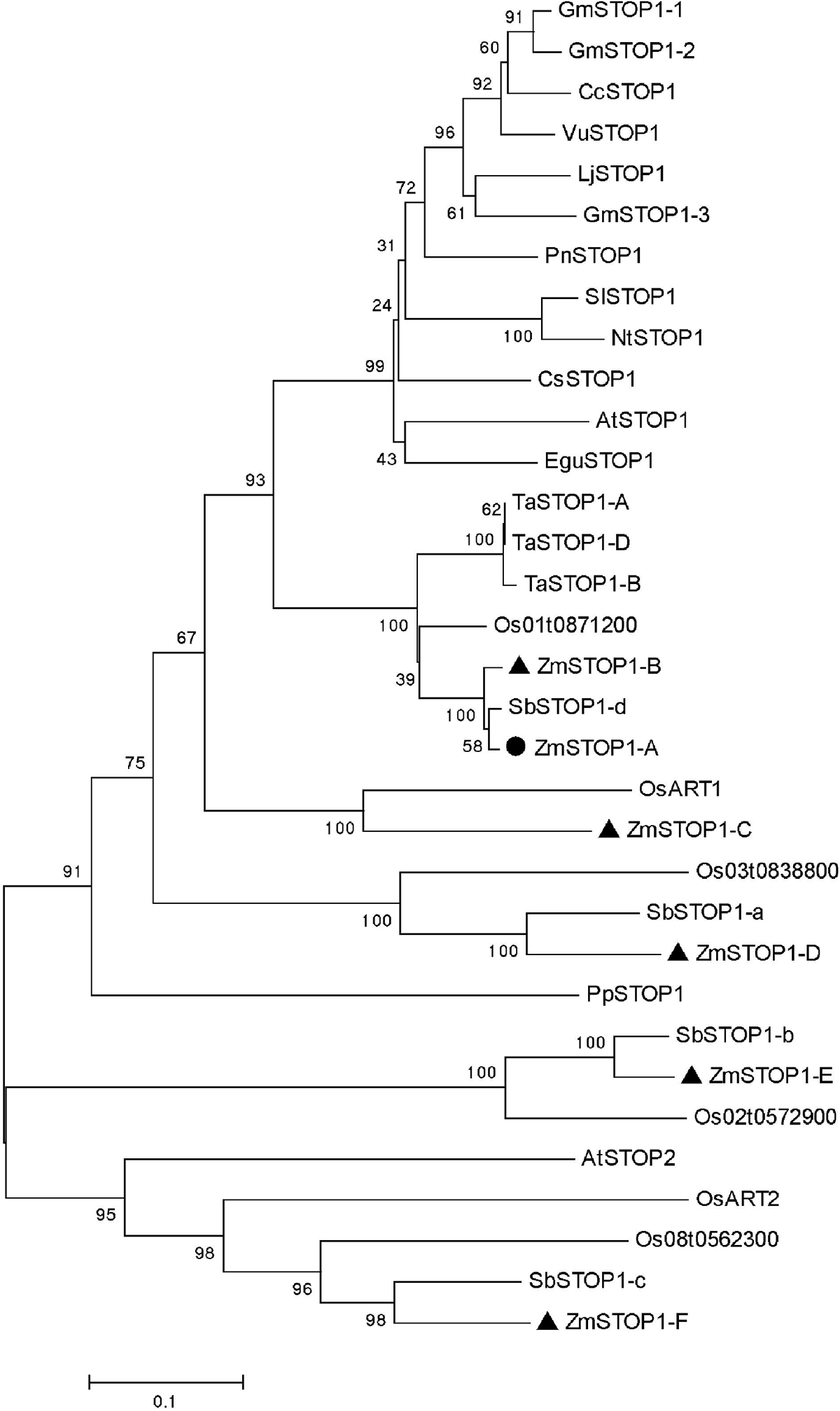
Phylogenetic tree of ZmSTOP1-like proteins. The phylogenetic tree was constructed based on an amino acid sequence alignment of STOP1 orthologs from several plant species.

### Expression pattern of *ZmSTOP1-A*

To examine whether these six *ZmSTOP1-like* genes were responsive to Al stress, their expression was analyzed by quantitative real-time PCR in maize seedling roots. As shown in **Figure S2**, *ZmSTOP1-like* gene expression was constitutive and not induced by Al toxicity. However, ZmSTOP1-A showed relatively high expression and was chosen for further Al-response analyses.

*ZmSTOP1-A* exhibited similar expression pattern in both roots and shoots with or without Al stress (**Figure 2A**). Further testing found that Al exposure did not affect *ZmSTOP1-A* expression in different root segments (0-5 mm, 5-10 mm, 10-20 mm) (**Figure 2B**), nor did low pH treatment(**Figure 2C**). Arabidopsis plants transformed with a *ZmSTOP1-A* promoter-GUS construct confirmed this constitutive expression pattern in roots, leaves and stems. These results demonstrate that *ZmSTOP1-A* is constitutively expressed in both roots and shoots, and not induced by Al toxicity and low pH (**Figure 2D, 2E**).

**Fig. 2.**
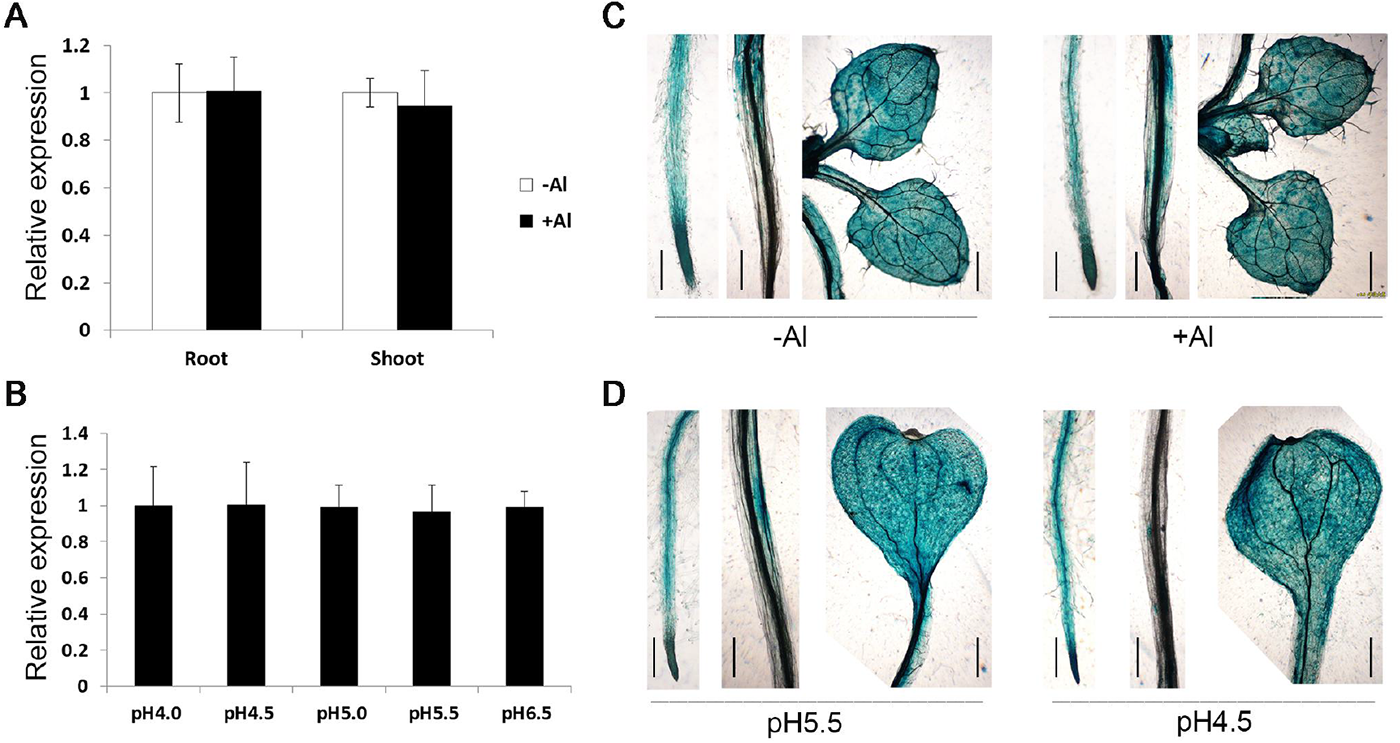
Spatial expression pattern of *ZmSTOP1-A* in maize seedlings. The expression of *ZmSTOP1-A* in different (A) organs, (B) root segments, and (C) under different pH. GUS staining of roots, stems and leaves of *promoter^ZmSTOP1-A^::GUS* transgenic *Arabidopsis* plants under (D) Al stress and (E) low pH. Al stress was performed in the solution containing 0 or 222μM Al [KAl(SO_4_) _2_] at pH 4.0 for 6 h. Mean values and SD (n = 3) are shown. Different letters indicate significant difference (Tukey’s test, *P* < 0.05). At least two independent transgenic lines were used for GUS staining. Bars = 200μm.

### Subcellular localization and transcriptional activation of ZmSTOP1-A

When transiently expressed in onion epidermal cells, ZmSTOP1-A::GFP fluorescence was only observed in the nucleus (**Figure 3A**) in comparison with 35S::GFP control which localized to both the nucleus and plasma membrane. This indicates ZmSTOP1-A is a nucleus protein.

**Fig. 3.**
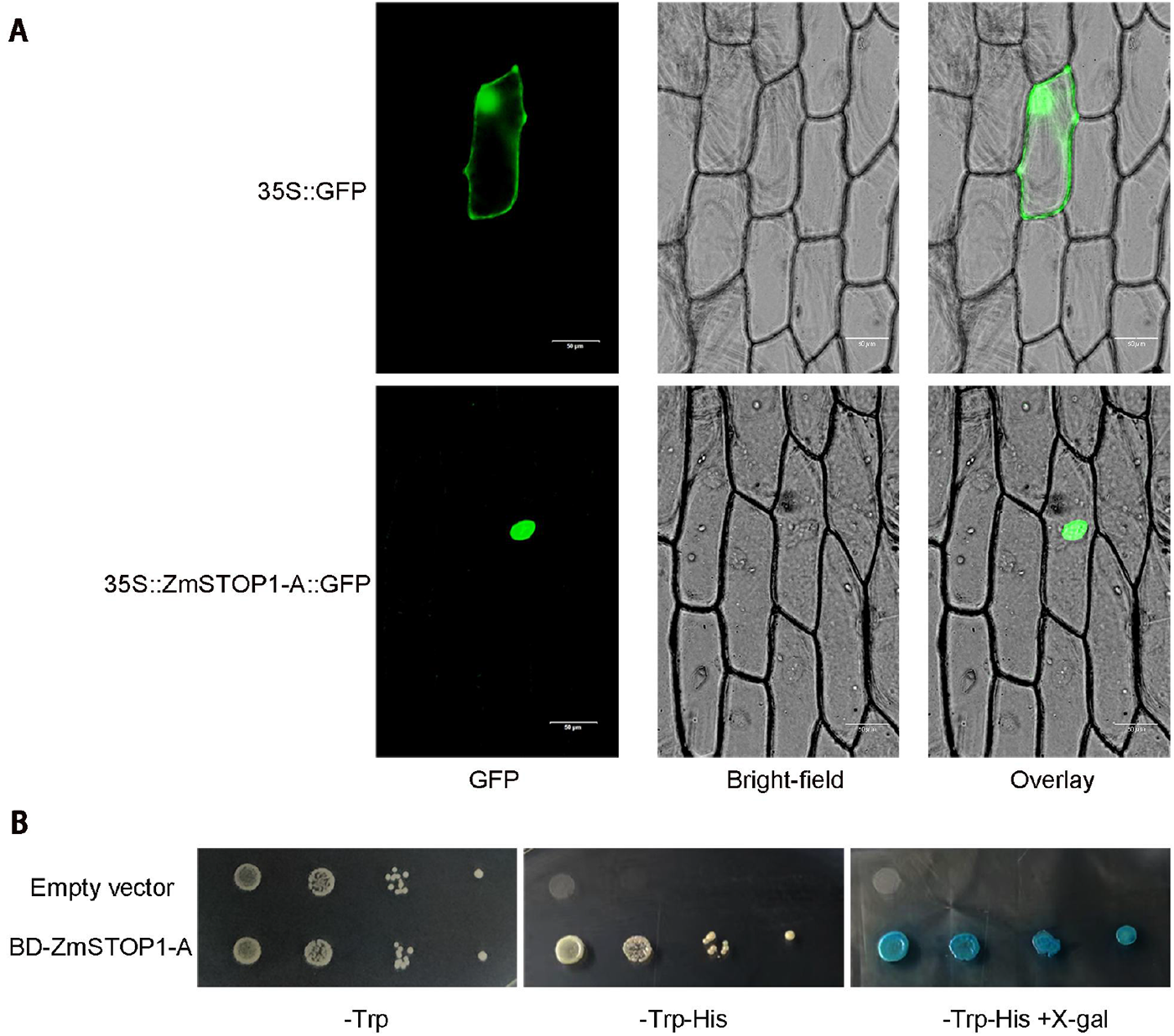
Subcellular localization and transactivation activity of ZmSTOP1-A. (A) *35S::GFP* (control) and *35S::ZmSTOP1-A::GFP* constructs were expressed in onion epidermal cells. Bars = 10μm. (B) *β*-galactosidase activity was indicated by blue color using X-gal as the substrate.

The transcriptional activation ability of ZmSTOP1-A was also verified in yeast system. As shown in Figure 3B, yeast cells transformed with empty vector p*GBKT7* did not grow on screening medium. However, yeast cells expressing p*GBKT7-ZmSTOP1-A* grew normally and turned blue after X-gal adding, indicating ZmSTOP1-A has transcriptional activation capability, a defining property of transcription factors.

### Ectopic expression of *ZmSTOP1-A* enhanced Al- and low pH-tolerance in transgenic Arabidopsis

Whether *ZmSTOP1-A* is also involved in Al toxicity and low pH stress in maize is unclear yet. To examine this, *ZmSTOP1-A* was transformed to *Atstop1* mutant and highly expressed transgenic lines *ZmSTOP1-A/Atstop1* #5 and #8 (**Figure S3**) were selected for genetic complementary evaluation. Under Al stress (pH 4.9+Al), the relative root growth (RRG) of *ZmSTOP1-A/Atstop1* #5 and #8 was slightly lower than wild-type (WT) but significantly higher than *Atstop1*, indicating that *ZmSTOP1-A* could partially restore the Al-induced root growth inhibition in *Atstop1* (**Figure 4A and 4B**). Similarly, at pH 4.9, *ZmSTOP1-A/Atstop1* (#5 and #8) fully restored RRG of *Atstop1* to WT levels (**Figure 4A and 4C**). These results demonstrate that *ZmSTOP1-A* facilitates Arabidopsis plants to antagonize both Al toxicity and low pH.

**Fig. 4.**
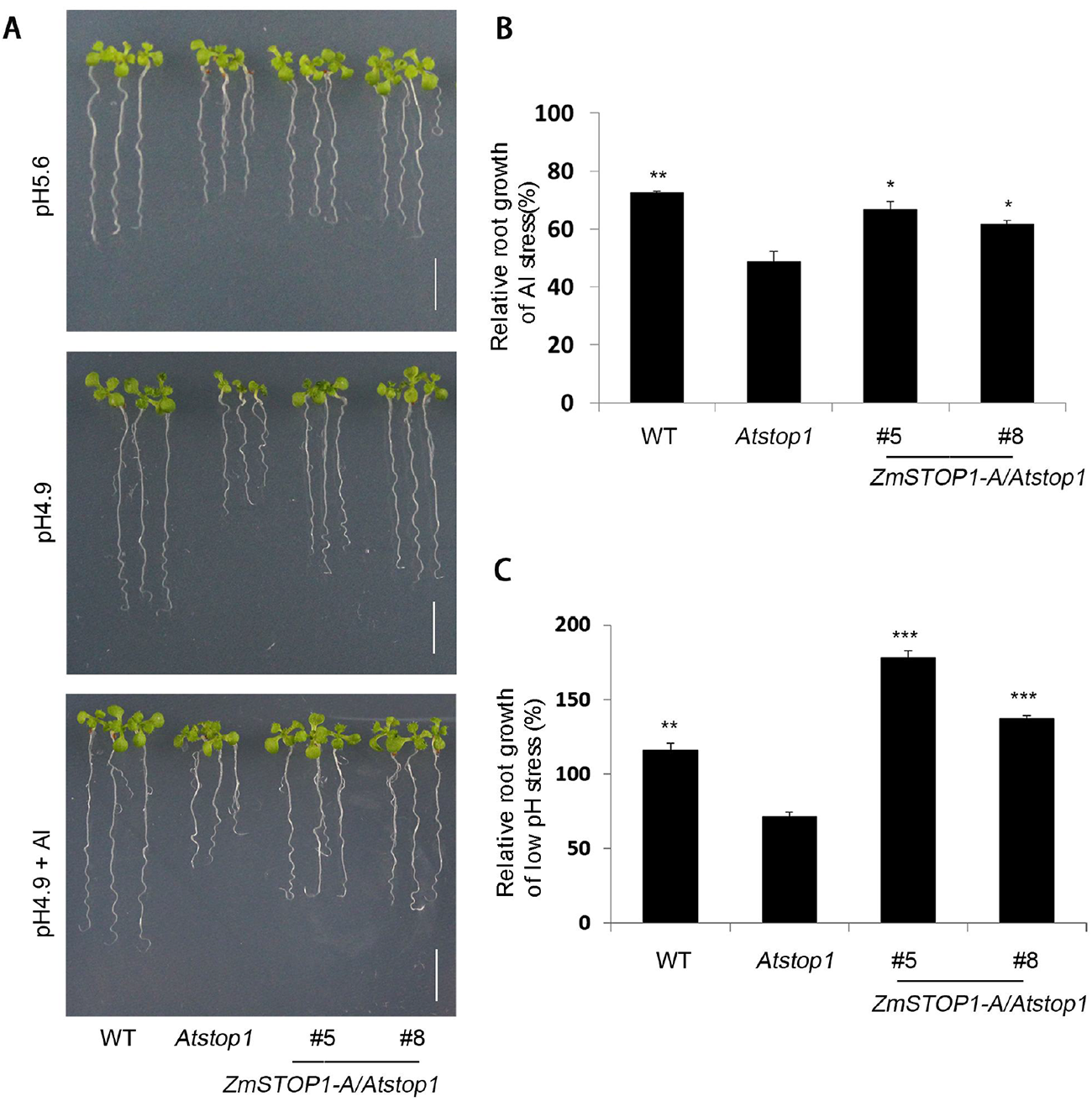
Phenotypes and relative root growth of *ZmSTOP1-A/Atstop1* under Al toxicity and low pH stress. (A)Phenotypes of WT, *Atstop1* and *ZmSTOP1-A/Atstop1*(#5 and #8) under 20μM AlCl_3_ (pH 4.9) and pH 4.9 conditions. Bars = 1 cm. (B) Relative root growth (RRG) of genetypes under 20μM AlCl3 (pH 4.9). (C) RRG of genotypes under pH 4.9. Means and SD (n = 10) are shown. Asterisks indicate significant differences compared to *Atstop1* (Tukey’s test; ∗ *P* < 0.05; ∗∗ *P* < 0.01; ∗∗∗ *P* < 0.001).

It has been shown that Arabidopsis AtSTOP1 responds to Al toxicity and low pH by regulating the expression of different set of genes (Sawaki et al., 2009). Thus, the expression of *AtALMT1*, *AtMATE* and *AtALS3* (Al-responsive), and *AtSTOP2*, *AtGDH1*, *AtGDH2*, *AtNRT1.1* and *AtCIPK23* (low pH-responsive) were examined here. Under Al stress, Al-responsive genes expression was partially restored in all *ZmSTOP1-A/Atstop1* plants compared to *Atstop1* (**Figure 5**). Under low pH conditions, *AtGDH1*, *AtGDH2* and *AtNRT1.1* but not *AtSTOP2* and *AtCIPK23* expression fully recovered in *ZmSTOP1-A/Atstop1* plants (**Figure 5**). These expression patterns align with the RRG of *ZmSTOP1-A/Atstop1* plants under Al toxicity and low pH conditions.

**Fig. 5.**
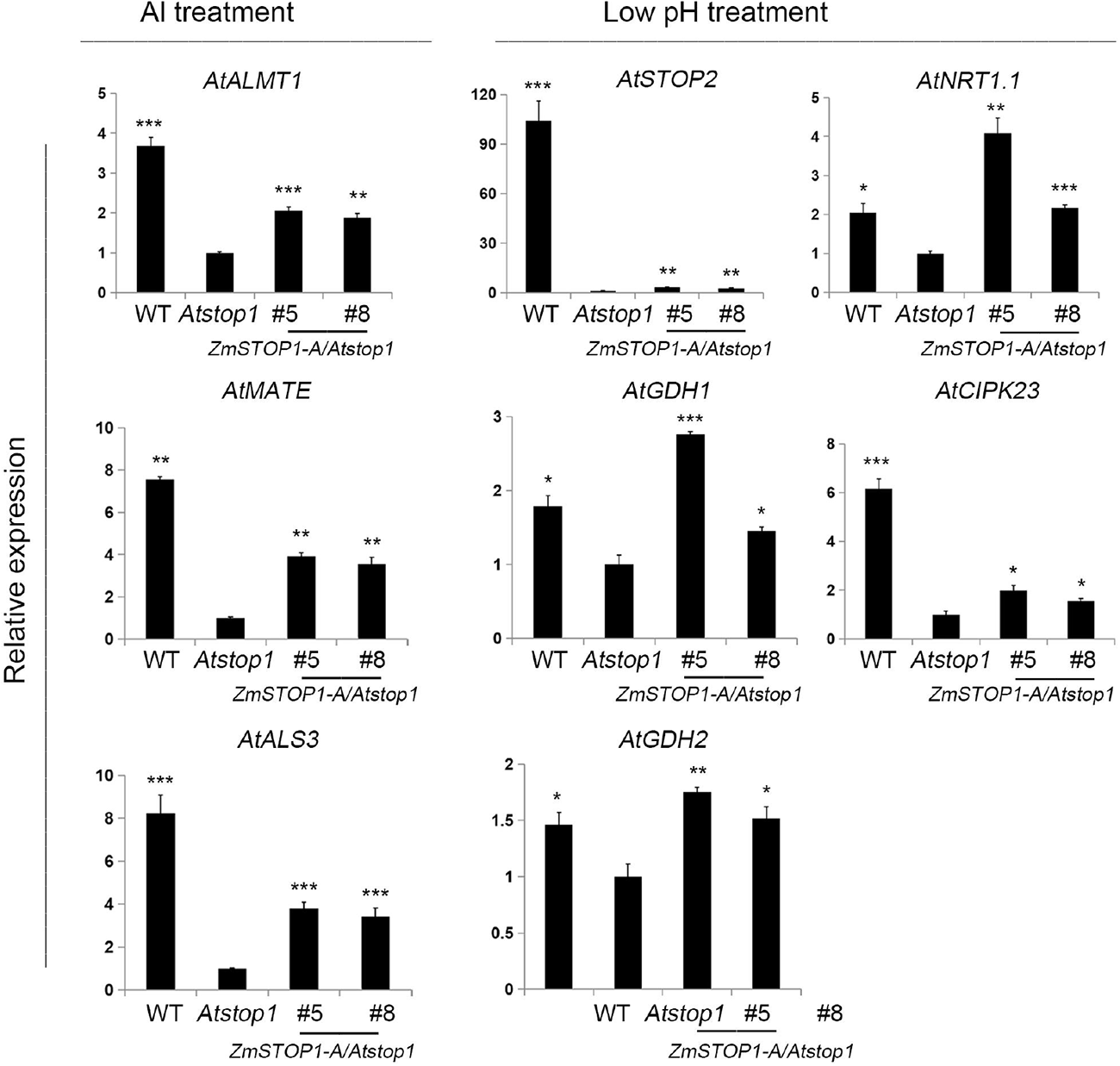
Expression of *ZmSTOP1-A* regulated genes under Al toxicity or low pH stress. WT, *Atstop1* and *ZmSTOP1-A/Atstop1* plants were exposed to 20μM AlCl_3_ (pH 4.9) or pH 4.9 for 24 h. Al-responsive genes *AtALMT1*, *AtMATE*, *AtALS3* and low pH-responsive genes *AtSTOP2*, *AtGDH1*, *AtGDH2*, *AtNRT1.1*, *AtCIPK23* were quantified by qRT-PCR using *AtACT2* as an internal control. Means values and SD (n = 3) are shown. Asterisks indicate significant differences compared to *Atstop1* (Tukey’s test; ∗ *P* < 0.05; ∗∗ *P* < 0.01; ∗∗∗ *P* < 0.001).

### *ZmSTOP1-A* partially rescued Al tolerance of *Atstop1* attributed to the secretion of organic acids

The primary mechanism for crops to detoxify Al is extrusion of organic acids from the root apex. As expected, under Al stress, the secretion of malate and citrate was significantly higher in *ZmSTOP1-A/Atstop1* plants compared to *Atstop1*, but still lower than WT (**Figure 6A and 6B**). This aligns with the expression of *AtALMT1* and *AtMATE*. Furthermore, Al content in *ZmSTOP1-A/Atstop1* roots was higher than WT but lower than *Atstop1* (**Figure 6D**). Hematoxylin staining confirmed these findings (**Figure 6C**). Together, these results indicate that organic acid extrusion is a key strategy by which ZmSTOP1-A alleviates Al toxicity of *Atstop1*.

**Fig. 6.**
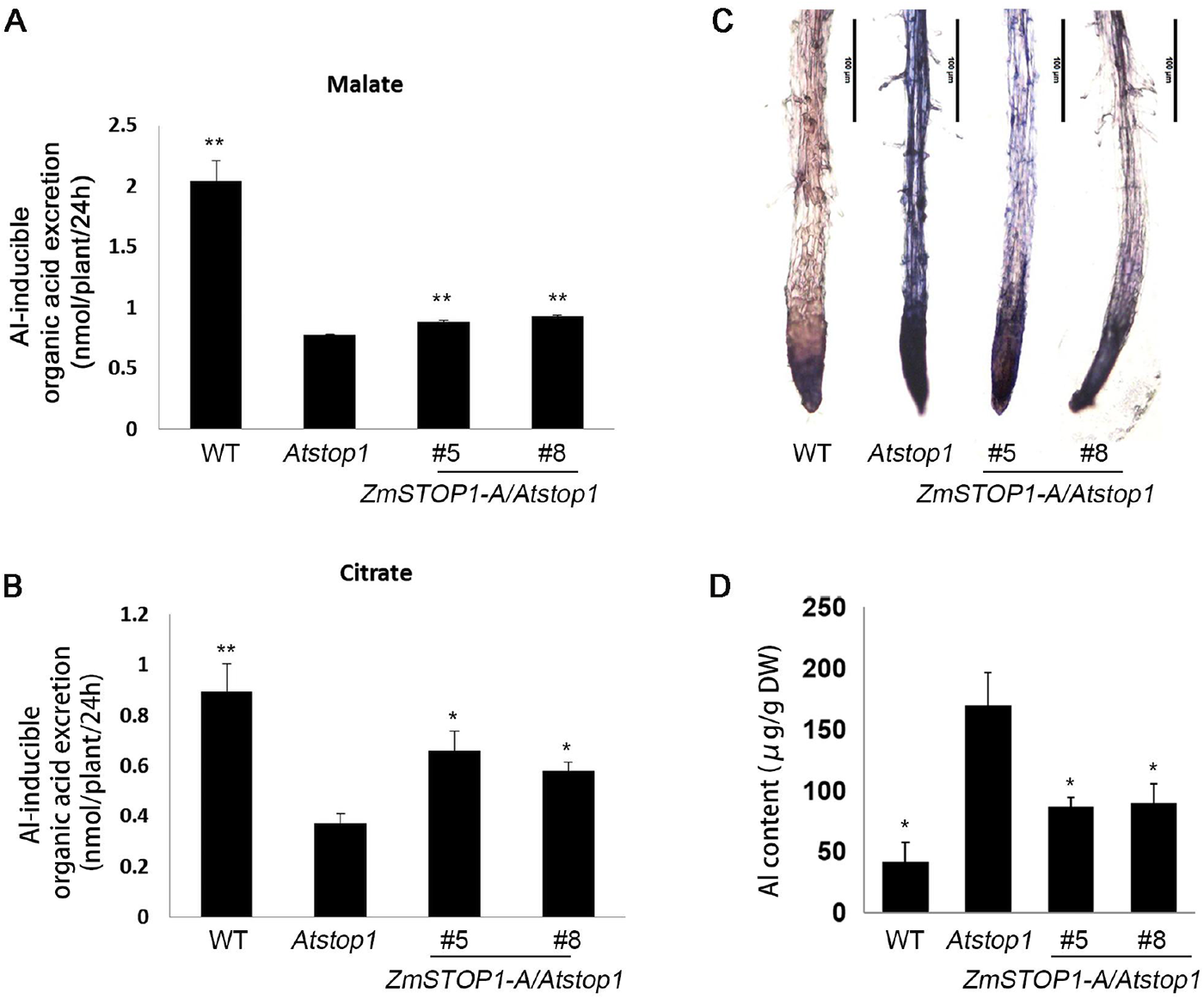
Overexpression of *ZmSTOP1-A* reduced Al accumulation in *Atstop1* mutant. Secretion of (A) citrate and (B) malate, (C) Al content, and (D) hematoxylin staining in roots of WT, *Atstop1* and *ZmSTOP1-A/Atstop1* plants after 24 h treatment with or without 20μM AlCl_3_ (pH 4.9). Values in A, B, and C represent mean ± SD (n ≥ 10). Asterisks indicate significant differences compared to *Atstop1* (Tukey’s test; ∗ *P* < 0.05; ∗∗ *P* < 0.01).

### *ZmSTOP1-A* responds to Al-induced oxidative stress in the *Atstop1* mutant

Al toxicity can induce ROS production, and scavenging of these ROS mitigates Al stress. Thus, the activities of antioxidant enzymes, SOD, CAT, POD and APX, which eliminate ROS, were examined. Under Al stress, O_2_^−^ accumulation was higher in all plant roots, but O_2_^−^ content in *ZmSTOP1-A/Atstop1* was similar to WT and lower than in *Atstop1* **(Figure 7A)**. In contrast, H_2_O_2_ increased in roots upon Al treatment, with no difference between +Al or -Al conditions **(Figure 7B)**. Meanwhile, SOD, CAT, POD and APX activities increased in Al-stressed roots. In *ZmSTOP1-A/Atstop1* plants, SOD, CAT and POD activities fully recovered to WT levels, except for APX which was similar across the tested plants **(Figure 7C-7F)**. The same pattern also occurred in shoots **(Figure S5).** This indicates that SOD, CAT, POD preferentially contribute to scavenging Al-induced ROS, particularly O_2_^−^.

**Fig. 7.**
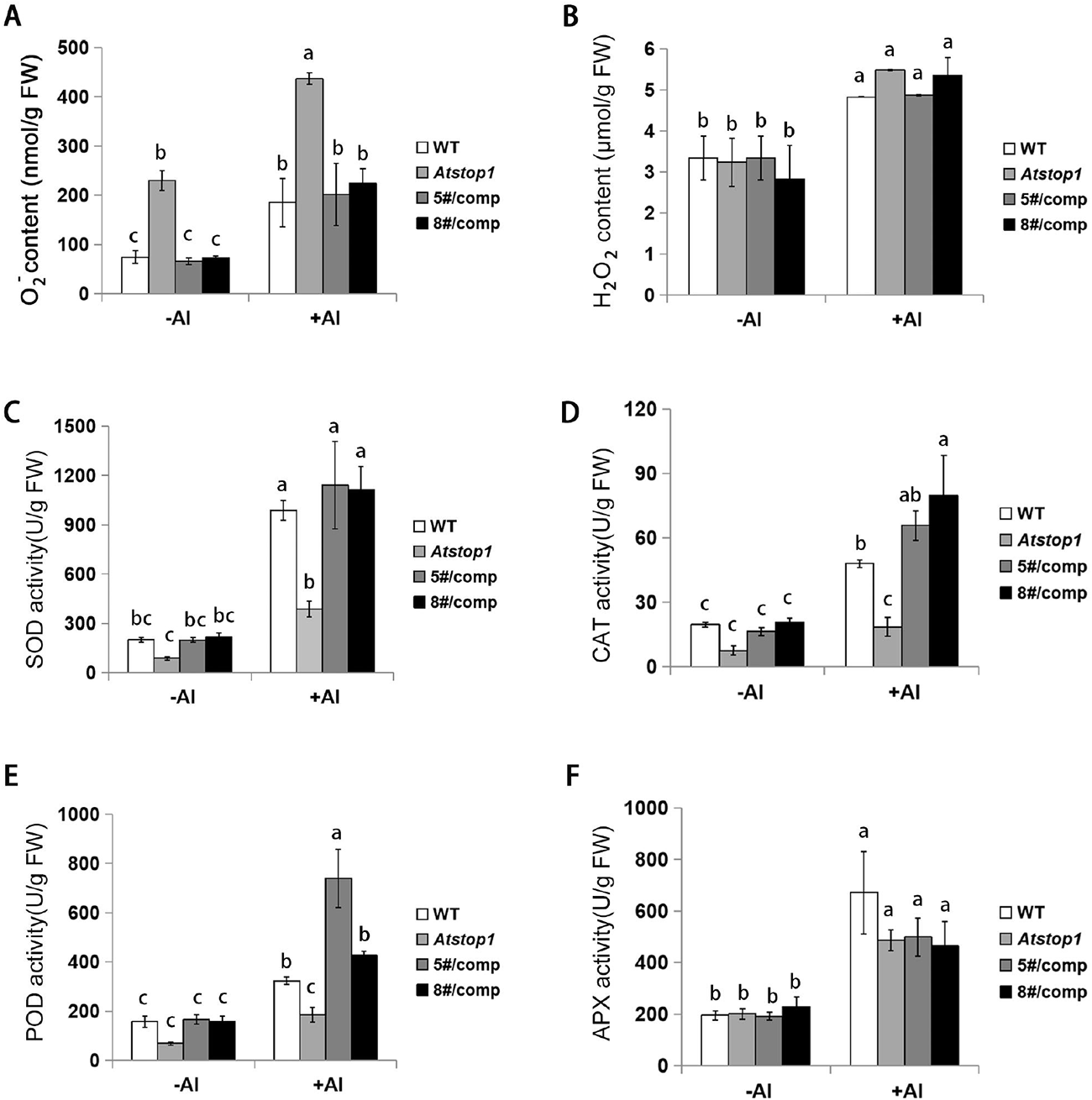
Effects of Al toxicity on reactive oxygen species in WT, *Atstop1*, and *ZmSTOP1-A/Atstop1* plants. Superoxide anion (O_2_^−^) (A) and hydrogen peroxide (H_2_O_2_) (B) content and activity of SOD (C), CAT (D), POD (E) and APX (F) in roots after 24 h exposure with 0 or 20μM AlCl_3_ (pH 4.9). Values are mean ± SD (n ≥ 10) of three separated experiments. Different letters indicate significant difference (Tukey’s test, *P* < 0.05).

### ZmSTOP1-A directly regulated *ZmMATE1* expression in maize

Organic acid extrusion is the primary Al tolerance mechanism in many plant species, with maize mainly secreting citrate via the important citrate transporter *ZmMATE1* (Maron et al., 2010). In order to examine if ZmSTOP1-A directly regulated *ZmMATE1*, a transient promoter assay was performed in maize protoplasts. Co-transformation of promoter*^ZmMATE1^:LUC* and *35S:ZmSTOP1-A* enhanced LUC signal compared to promoter*^ZmMATE1^:LUC* alone (**Figure 8B**). This demonstrates that ZmSTOP1-A can activate *ZmMATE1* transcription.

**Fig. 8.**
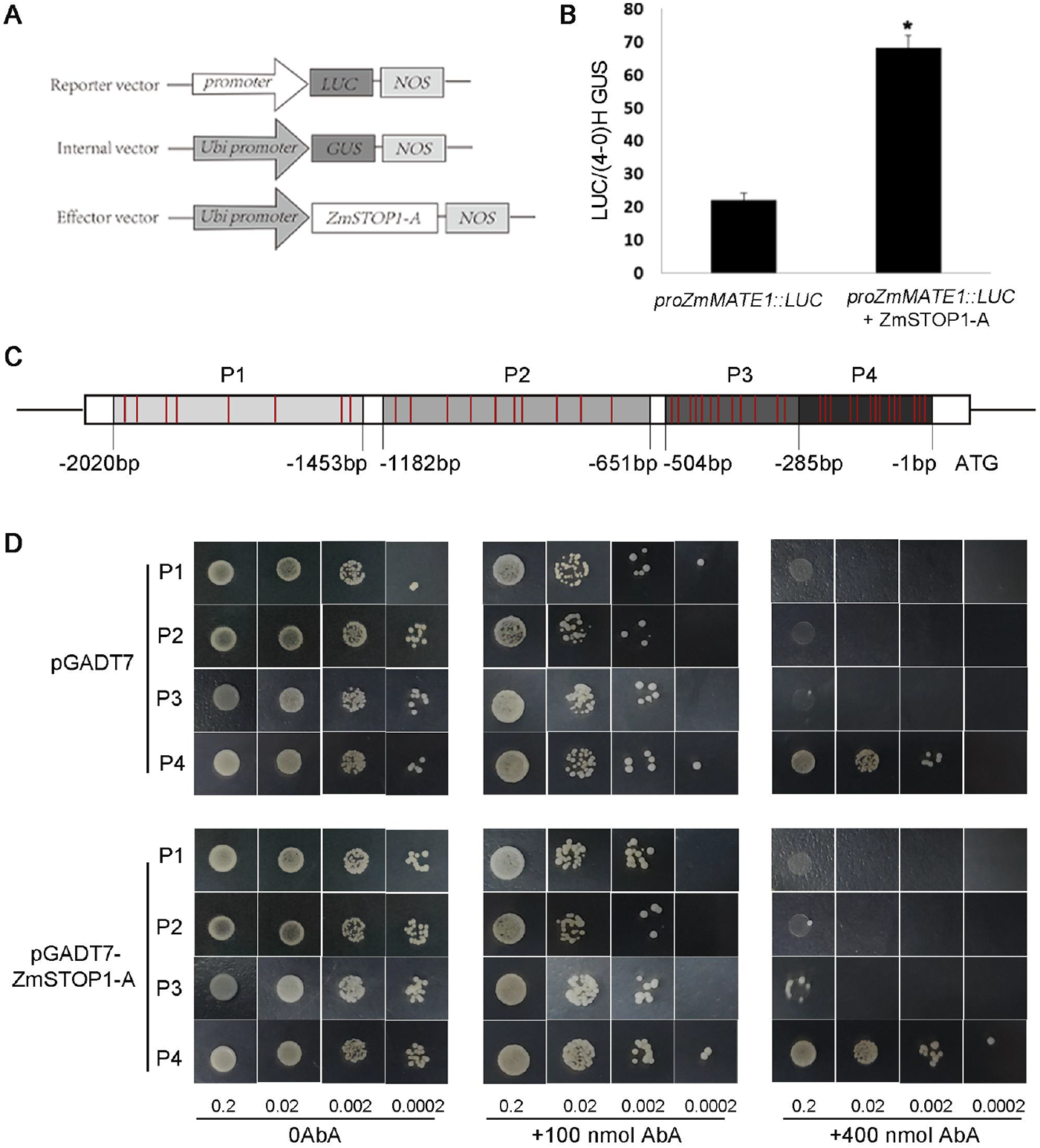
*ZmSTOP1-A* activated *ZmMATE1* expression by binding to its promoter fragments. (A) Schematic of transient expression vectors. (B) ZmSTOP1-A activation of *ZmMATE1* in maize protoplast assay. Values are means ± SD (n = 3). Asterisks indicate significant differences between with and without effector vectors (Tukey’s test, *P* < 0.05). (C) Schematic diagram of *ZmMATE1* promoter bait fragments (P1-P4) used to construct reporter vectors. Red indicates GGNVS *cis*-elements. (D) Binding of ZmSTOP1-A to different *ZmMATE1* promoter fragments in yeast one-hybrid assay.

In addition, the *ZmMATE1* promoter was truncated into P1-P4 four fragments for yeast one-hybrid analysis. P1 (-2020 to -1453), P2 (-1182 to -651), P3 (-504 to -285), and P4 (-285 to -1) contain 8, 10, 12, and 14 Al-responsive GGNVS *cis*-elements respectively (**Figure 8C**). All yeast grew normally at 100 ng/ml AbA, but were inhibited at 400ng/ml AbA. Compared to the pGADT7 control, yeast with *ZmSTOP1-A* promoter fragment grew much better, declining in rank P4, P3, P2, P1 (**Figure 8D**). These demonstrate that ZmSTOP1-A directly interacts with *ZmMATE1* promoter, with binding strength positively correlating with the GGNVS *cis*-element number.

## Discussion

### *ZmSTOP1-A* is involved in both Al stress and low pH stress responses

Al toxicity is the major limiting factor for plant growth in acid soils. Identification of Al-response genes and elucidating their roles is essential to understand the molecular mechanisms of Al toxicity, and to develop molecular-assistant breeding approaches to improve crop yields under Al stress. The transcription factor AtSTOP1 and its rice homolog OsART1 are master regulators of over twenty Al-responsive genes in Arabidopsis and rice, respectively (Iuchi et al., 2007; Yamaji et al., 2009). STOP1-like transcription factors have now been found across many plant species, where they are involved in both Al stress and low pH stress responses, unless not yet investigated (Garcia-Oliveira et al., 2013; Ohyama et al., 2013; Sawaki et al., 2014). For example, AtSTOP1 was first identified in Arabidopsis from a proton-sensitive mutant, and later shown to mediate Al tolerance (Iuchi et al., 2007). Similarly, GmSTOP1-1 and -3 in soybean (Wu et al., 2018), ScSTOP1 in rye (Silva-Navas et al., 2021), PpSTOP1 in moss, PnSTOP1 in poplar, LjSTOP1 in lotus and NtSTOP1 in tobacco (Ohyama et al., 2013), had all been shown to function in both Al and low pH stresses by complementing the Arabidopsis *Atstop1* mutant. In this study, *ZmSTOP1-A/Atstop1* plants also fully or partially restored the WT phenotype via regulating distinct Al- and low pH-responsive genes (**Figure 4 and 5**), consistent with finding on other STOP1-like transcription factors using this complementary system.

Nevertheless, the downstream genes regulated by STOP1-like proteins differ somewhat between species in response to both Al and low pH stresses. For example, *ZmSTOP1-A* rescued low pH-sensitivity of *Atstop1* by upregulating *AtSTOP2*, *AtGDH1*, *AtGDH2*, *AtCIPK23* and *AtNRT1.1* (**Figure 5**), while *NtSTOP1*, *PpSTOP1*, *EguSTOP1* etc. act through *AtSTOP2*, *AtPGIP1*, *AtCIPK23* (Ohyama et al., 2013; Sawaki et al., 2014) . *VuSTOP1* mediates via *AtSTOP2*, *AtGDH1*, *AtPGIP1*, and *AtCIPK23* (Fan et al., 2015). Beyond that, STOP1-like gene expression patterns vary. Some are Al-inducible, like *OsART2*, *VuSTOP1*, *TaSTOP1-A*, *SbSTOP1s, GmSTOP1-1* (Garcia-Oliveira et al., 2013; Fan et al., 2015; Che et al., 2018; Huang et al., 2018; Wu et al., 2018), while others including *TaSTOP1-B*/-*D, AtSTOP1, AtSTOP2*, *EguSTOP1, HvATF1* (Al-tolerance transcription factor) and *OsART1* are constitutively expressed regardless of Al exposure (Iuchi et al., 2007; Yamaji et al., 2009; Garcia-Oliveira et al., 2013; Kobayashi et al., 2014; Sawaki et al., 2014; Wu et al., 2020). Similar variation exists under low pH (Table S2). But in fact, post-transcriptional modifications including SUMOylation, ubiquitination and phosphorylation, can modulate the stabilization of constitutively expressed STOP1-like protein, as demonstrated by AtSTOP1. These findings reveal that STOP1-like proteins have developed divergent function for plants species to adapt to fluctuating environment in acid soil.

In this study, *ZmSTOP1-A* expression was not affected by Al toxicity and low pH (**Figure 2**), similar to STOP1-like genes such as *AtSTOP1* and *OsART1* (Iuchi et al., 2007; Yamaji et al., 2009). Considering the diverse post-translational regulation of AtSTOP1, such modifications likely also modulate ZmSTOP1-A. Future work could examine potential ZmSTOP1-A modification under Al stress by fusing fluorescent reporters or epitope tags for genetic and biochemical detection.

### *ZmSTOP1-A* enhanced Al tolerance in Arabidopsis mainly through external extrusion mechanism

Ectopic expression in the Arabidopsis *Atstop1* mutant is a widely used system to assay Al tolerance genes. For instance, under Al stress condition, *VuSTOP1* and *AtSTOP2* partially rescued *Atstop1* Al sensitivity by restoring the expression of *AtALS3* and *AtMATE*, but not *AtALMT1* (Kobayashi et al., 2014; Fan et al., 2015). Similarly, *PpSTOP1* partially recovered *AtALMT1*, *AtMATE* and *AtALS3* expression and roots growth (Ohyama et al., 2013). Here, *ZmSTOP1-A* also partially restored *AtALMT1*, *AtMATE* and *AtALS3* expression in *Atstop1* under Al stress (**Figure 5**), correlating with increased organic acid secretion and lower Al accumulation (**Figure 6**). This consists with findings for *NtSTOP1* (Ohyama et al., 2013), suggesting the STOP1-organic acid transporter module is highly conserved for Al detoxification across plants.

It is notable that *STOP1-like* genes in most species are predominant expressed in roots with very low shoot levels, such as *OsART1*/*2* in rice (Yamaji et al., 2009; Che et al., 2018), *VuSTOP1* in rice beans (Fan et al., 2015), *SbSTOP1S* in sweet sorghum (Huang et al., 2018), and *GmSTOP1s* in soybean (Wu et al., 2018). In contrast, *ZmSTOP1-A* was expressed almost equally in roots and shoots of maize seedlings, implying a potential role for shoots in maize Al detoxification. *HvATF1* in barley follows this expression pattern, but its Al response was not explored (Wu et al., 2020). In *ZmSTOP1-A/Atstop1* plants here, both the internal tolerance gene *AtALS3* and external exclusion genes *AtALMT1* and *AtMATE* were upregulated (**Figure 5**). Meanwhile, Al content in *ZmSTOP1-A/Atstop1* roots and shoots sharply declined to WT level as in comparison with *Atstop1* (**Figure 6 and Figure S4**). This reveals external exclusion dominates in *ZmSTOP1-A/Atstop1* plants. The low Al absorption and root–to-shoot transport was insufficient to impact shoot Al content. This suggests *ZmSTOP1-A* may confer Al tolerance in maize primary by regulating organic acid transporter expression, as demonstrated by other STOP1 transcription factors. For example, *AtSTOP1* regulates *AtALMT1* and *AtMATE* for organic acid secretion (Iuchi et al., 2007; Sawaki et al., 2009); *OsART1* mediates citrate secretion by controlling *OsFRDL4* and *OsFRDL2* expression (Yamaji et al., 2009), and *CcSTOP1* directly regulates *CcMATE1* expression (Daspute et al., 2018).

Additionally, *ZmMATE1* and *ZmMATE6* were previously reported to confer maize Al tolerance through citrate release (Maron et al., 2010; Du et al., 2021). Here, the transient assay in maize protoplasts and yeast one-hybrid analysis verified that ZmSTOP1-A directly binds to the Al-responsive GGNVS *cis*-element to activate *ZmMATE1* and detoxify Al (**Figure 8B**). Interestingly, ZmSTOP1-A activation of *ZmMATE1* positively correlated with the GGNVS *cis*-element number in the *ZmMATE1* promoter fragment (**Figure 8D**). This is consistent with findings for other organic transporters like HLALMT1 and OsFDRL4, which are activated by STOP1 proteins and show higher expression in Al-tolerant cultivars with more GGNVS elements (Chen et al., 2013; Yokosho et al., 2016). Therefore, increase GGNVS elements in promoters of organic acid transporter genes could be a strategy for breeding Al-tolerant crops.

### *ZmSTOP1-A* eliminated the Al-induced oxidative stress in *Atstop1*

Al toxicity triggers oxidative stress in many plants. Jay et al. found that Al toxicity caused excessive ROS (H_2_O_2_ and O_2_^−^) accumulation in rice roots, activating antioxidant enzymes including SOD, APX, POX, GR, CAT, DHAR, MDHAR. Al-resistant varieties displayed stronger antioxidant defences than sensitive ones (Awasthi et al., 2019). Additionally, overexpression of the Arabidopsis peroxidase gene *AtPrx64* improved tobacco Al tolerance by scavenging ROS and reducing root Al accumulation (Wu et al., 2017). Recently, *ZmAT6* overexpression in maize enhanced Al tolerance by dramatically increasing SOD and POD antioxidant enzyme activities (Du et al., 2020). In this study, *ZmSTOP1-A* also increased SOD, POD and CAT activities in *Atstop1* roots and shoots, coordinately with the eliminated ROS (**Figure 7 and FigureS5**). This demonstrates ROS scavenging assists Al detoxification whether directly or indirectly.

In summary, ZmSTOP1-A, a C2H2 zinc finger transcription factor involved in Al toxicity and low pH responses, was constitutively and equally expressed in maize roots and shoots. ZmSTOP1-A conferred tolerance to both Al toxicity and low pH stress. Regarding Al toxicity, *ZmSTOP1-A* increased organic acid secretion and reduced Al accumulation by upregulating *AtALMT1* and *AtMATE* expression. It also enhanced antioxidant content to detoxify Al. In addition, *ZmMATE1* is directly activated the maize organic acid transporter ZmSTOP1-A in, a manner dependent on the GGNVS element. Overall, these results demonstrate that ZmSTOP1-A plays an important role in responding to Al toxicity and low pH by regulating organic acid secretion and ROS scavenging.

## Materials and Methods

### Plant materials and culture conditions

The Al-tolerant maize inbred line 178 was used in this study. Maize seedling were cultivated in nutrient solution under growth conditions as previously described (Maron et al., 2010; Ligaba et al., 2012) . After 24 h pretreatment, two-leaf stage seedlings were exposed to the same solution with or without 222μM Al [KAl(SO_4_) _2_] at different pH (4.0, 4.5, 5.0, 5.5, 6.0 and 6.5) for 6 h. All treatments were performed in three replicates for RNA extraction (Iuchi et al., 2007).

Wild-type (WT) Col-0, *Atstop1* mutant (At1g34370, SALK_114108) and transgenic Arabidopsis were grown under condition as previously described by Kobayashi et al (Iuchi et al., 2007).

### Phylogenetic tree construction

Based on the sequence homolgy to proteins of AtSTOP1 and OsART1, six homologs, ZmSTOP1-A(Zm00001d042686), ZmSTOP1-B(Zm00001d012260), ZmSTOP1-C(Zm00001d023558), ZmSTOP1-D(Zm00001d034783), ZmSTOP1-E(Zm00001d016911) and ZmSTOP1-F(Zm00001d031895), were identified in maize. A phylogenetic tree of STOP1-like proteins from maize and other species was constructed using the neighbor-joining method in MEGA **5.1.**

### RNA extraction and quantitative real-time RT-PCR

Total RNA extraction and quantitative RT-PCR were performed as previously described (Du et al., 2020). All primers were designed using Primer 5.0 software (Table S1). *ZmGAPDH* and *AtACT2* were used as reference genes. Each sample was assayed in at least three technical replicates.

### Subcellular localization

The full-length ORF of *ZmSTOP1-A*, excluding the stop codon, was fused to *GFP* in pCAMBIA2300 to generate the *ZmSTOP1-A::GFP* fusion construct. This *ZmSTOP1-A::GFP* vector and a *35S::GFP* control vector were transiently expressed in onion epidermal cells as described previously by Li et al (Li et al., 2021).

### GUS staining assay

A 2.1 kb region upstream of the start codon of *ZmSTOP1-A* was amplified and cloned into vector pCAMBIA3301. GUS staining was performed as previous described (Jefferson, 1987).

### Determination of organic acids secretion and Al content

After Al stress treatment, root exudate organic acids concentrations in Arabidopsis were measured and hematoxylin staining performed as previously described (Du et al., 2021). Root and shoot samples were collected separately and Al content qualified by inductively coupled plasma mass spectrometry (ICP-MS, PerkinElmer, NexlON 2000).

### Transient expression assay in maize protoplasts

As described by Li et al. (Li et al., 2021), the reporter vector was generated by cloning the *ZmMATE1* promoter into the pBI221 vector. The effector vector was created by inserting the coding sequences of *ZmSTOP1-A*, driven by an ubiquitin promoter, into the *Pst*Ⅰand *Bam*HⅠ sites of pBI221. The ubi::*GUS* was used as the internal vector. After 12-14 hours culturing in the dark, the co-transformed maize 178 protoplasts containing the reporter vector, effecter vector, and internal vector (at a 2:1:1 molar ratio) were observed for LUC and GUS signal. The experiments were conducted with three independent replicates.

### Transcriptional activity detection and yeast one-hybrid assay

For the yeast transactivation assay, the bait vector pGBKT7 with *ZmSTOP1-A* fused to the GAL4 DNA-binding domain (BD) was used to transform the Y2H gold yeast strain as previously described (Huang et al., 2018).

For the yeast one-hybrid assay (Lou et al., 2020), the promoter region of *ZmMATE1* was amplified and cloned into the pAbAi vector. *ZmSTOP1-A* was fused to the *GAL4* activation domain (AD) in the pGADT7 vector. Finally, this pair of constructs was then introduced into Y1H gold yeast strain.

### Determination of ROS content and antioxidant enzyme activity

O^2−^ and H_2_O_2_ content were individually determined following the method of Erich et al (Erich et al., 1975). Activities of SOD (EC 1.15.1.1), CAT (EC 1.11.1.6), APX (EC 1.11.1.11) and POX (EC 1.11.1.7) were carried out as described (Awasthi et al., 2019).

## Acknowledgements

This work was supported by the National Natural Science Foundation of China (No. 3 2171951) and Program for International Science and Technology Cooperation Project s of Chengdu (2023-GH02-00040-HZ).

## Author contributions

Suzhi Zhang designed the experiment and conception; Chan Liu performed the experiments and wrote the manuscript; Xiaoqi Hu, Lei Zang, Xiaofeng Liu, Yuhui Wei, Xue Wang, Xinwu Jin, Chengfeng Du and Yan Yu helped to collect and analyze the data; Wenzhu He provided the valuable suggestions on the manuscript; Suzhi Zhang obtained the funding and were responsible for this manuscript. All authors read and approved the manuscript.

## Supplemental data

The following materials are available in the online version of this article.

Supplemental Figure S1. Sequence alignment of ZmSTOP1-A with other STOP1-like transcription factors.

Supplemental Figure S2. Relative expression of *ZmSTOP1-like* gene in maize seedling roots with 0 or 222 μM Al [KAl(SO_4_) _2_].

Supplemental Figure S3. Screening of *Atstop1* mutant, *ZmSTOP1-A/Atstop1* plants and *ZmSTOP1-A* expression in *Atstop1*.

Supplemental Figure S4. Al contents in shoots of WT, *Atstop1* and *ZmSTOP1-A/Atstop1* plants after 24 h exposure in 20 μM AlCl_3_ (pH 4.9).

Supplemental Figure S5. Effects of Al toxicity on reactive oxygen species in WT, *Atstop1*, and *ZmSTOP1-A/Atstop1* shoots.

Supplemental Table S1. The primer used in this study.

Supplemental Table S2. Summary of STOP1-like genes involved in Al and/or low pH tolerance.

## Data availability

The datasets presented in this study can be found in online repositories. The names of the repository/repositories and accession number(s) can be found in the article/Supplementary Material.

## References

Awasthi JP, Saha B, Panigrahi J, Yanase E, Koyama H, Panda SK (2019) Redox balance, metabolic fingerprint and physiological characterization in contrasting North East Indian rice for Aluminum stress tolerance. Scientific reports 9: 8681

Che J, Tsutsui T, Yokosho K, Yamaji N, Ma JF (2018) Functional characterization of an aluminum (Al)-inducible transcription factor, ART2, revealed a different pathway for Al tolerance in rice. The New phytologist 220: 209–218

Chen ZC, Yokosho K, Kashino M, Zhao F, Yamaji N, Ma JF (2013) Adaptation to acidic soil is achieved by increased numbers of cis-acting elements regulating ALMT1 expression in Holcus lanatus. The Plant Journal: n/a-n/a

Daspute AA, Kobayashi Y, Panda SK, Fakrudin B, Kobayashi Y, Tokizawa M, Iuchi S, Choudhary AK, Yamamoto YY, Koyama H (2018) Characterization of CcSTOP1; a C2H2-type transcription factor regulates Al tolerance gene in pigeonpea. Planta 247: 201–214

Du H, Hu X, Yang W, Hu W, Yan W, Li Y, He W, Cao M, Zhang X, Luo B, Gao S, Zhang S (2021) ZmXTH, a xyloglucan endotransglucosylase/hydrolase gene of maize, conferred aluminum tolerance in Arabidopsis. Journal of Plant Physiology 266: 153520

Du H, Huang Y, Qu M, Li Y, Hu X, Yang W, Li H, He W, Ding J, Liu C (2020) A Maize ZmAT6 gene confers aluminum tolerance via reactive oxygen species scavenging. Frontiers in plant science 11: 1016

Du H, Liu C, Jin X, Du C, Yu Y, Luo S, He W, Zhang S (2022) Overexpression of the aldehyde dehydrogenase gene ZmALDH confers aluminum tolerance in Arabidopsis thaliana. International Journal of Molecular Sciences 23: 477

Du H, Ryan PR, Liu C, Li H, Hu W, Yan W, Huang Y, He W, Luo B, Zhang X, Gao S, Zhou S, Zhang S (2021) ZmMATE6 from maize encodes a citrate transporter that enhances aluminum tolerance in transgenic Arabidopsis thaliana. Plant science : an international journal of experimental plant biology 311: 111016

Erich E, Claus S, Adelheid H (1975) Determination of Superoxide Free Radical Ion and Hydrogen Peroxide as Products of Photosynthetic Oxygen Reduction. Zeitschrift für Naturforschung C 30: 53–57

Ezaki B, Suzuki M, Motoda H, Kawamura M, Nakashima S, Matsumoto H (2004) Mechanism of gene expression of Arabidopsis glutathione S-transferase, AtGST1, and AtGST11 in response to aluminum stress. Plant Physiology 134: 1672–1682

Fan W, Lou HQ, Gong YL, Liu MY, Cao MJ, Liu Y, Yang JL, Zheng SJ (2015) Characterization of an inducible C2 H2 -type zinc finger transcription factor VuSTOP1 in rice bean (Vigna umbellata) reveals differential regulation between low pH and aluminum tolerance mechanisms. The New phytologist 208: 456–468

Fang Q, Zhang J, Yang DL, Huang CF (2021) Arabidopsis thalianaThe SUMO E3 ligase SIZ1 partially regulates STOP1 SUMOylation and stability in. Plant signaling & behavior 16: 1899487

Garcia-Oliveira AL, Benito C, Prieto P, de Andrade Menezes R, Rodrigues-Pousada C, Guedes-Pinto H, Martins-Lopes P (2013) Molecular characterization of TaSTOP1 homoeologues and their response to aluminium and proton (H+) toxicity in bread wheat (Triticum aestivum L.). BMC plant biology 13: 1–13

Guo J, Zhang Y, Gao H, Li S, Wang Z, Huang C (2020) Mutation ofHPR1encoding a component of the THO/TREX complex reduces STOP1 accumulation and aluminium resistance inArabidopsis thaliana. The New Phytologist 228

Huang S, Gao J, You J, Liang Y, Guan K, Yan S, Zhan M, Yang Z (2018) Sorghum bicolorIdentification of STOP1-Like Proteins Associated With Aluminum Tolerance in Sweet Sorghum (L.). Frontiers in plant science 9: 258

Iuchi S, Koyama H, Iuchi A, Kobayashi Y, Kitabayashi S, Kobayashi Y, Ikka T, Hirayama T, Shinozaki K, Kobayashi M (2007) Zinc finger protein STOP1 is critical for proton tolerance in Arabidopsis and coregulates a key gene in aluminum tolerance. Proceedings of the National Academy of Sciences of the United States of America 104: 9900–9905

Jefferson RA (1987) GUS fusions: beta-glucuronidase as a sensitive and versatile gene fusion marker in higher plants. Embo Journal 6: 3901–3907

Kobayashi T, Nishizawa NK (2012) Iron uptake, translocation, and regulation in higher plants. Annual review of plant biology 63: 131–152

Kobayashi Y, Ohyama Y, Kobayashi Y, Ito H, Iuchi S, Fujita M, Zhao CR, Tanveer T, Ganesan M, Kobayashi M, Koyama H (2014) STOP2 activates transcription of several genes for Al- and low pH-tolerance that are regulated by STOP1 in Arabidopsis. Molecular plant 7: 311–322

Kochian LV, Hoekenga OA, Pi Eros MA (2004) HOW DO CROP PLANTS TOLERATE ACID SOILS? MECHANISMS OF ALUMINUM TOLERANCE AND PHOSPHOROUS EFFICIENCY. Annual Review of Plant Biology 55: 459–493

Kochian LV, Pi Eros MA, Liu J, Magalhaes JV (2015) Plant Adaptation to Acid Soils: The Molecular Basis for Crop Aluminum Resistance. Annual Review of Plant Biology 66: 571–598

Li H, Wang Y, Xiao Q, Luo L, Zhang C, Mao C, Du J, Long T, Cao Y, Yi Q, Wang Y, Li Y, Huang H, Liu H, Hu Y, Yu G, Liu Y, Zhang J, Huang Y (2021) Transcription factor ZmPLATZ2 positively regulate the starch synthesis in maize. Plant Growth Regulation 93: 291–302

Lidon FC, Barreiro MDG (2002) An overview into aluminum toxicity in maize. Bulg. J. Plant Physiol 28: 96–112

Ligaba A, Maron L, Shaff J, Kochian L, Pineros M (2012) Maize ZmALMT2 is a root anion transporter that mediates constitutive root malate efflux. Plant, cell & environment 35: 1185–1200

Lou HQ, Fan W, Jin JF, Xu JM, Chen WW, Yang JL, Zheng SJ (2020) A NAC-type transcription factor confers aluminium resistance by regulating cell wall-associated receptor kinase 1 and cell wall pectin. Plant, cell & environment 43: 463–478

Ma JF, Ryan PR, Delhaize E (2001) Aluminium tolerance in plants and the complexing role of organic acids. Trends in plant science 6: 273–278

Magalhaes JV, Liu J, Guimarães CT, Lana UGP, Alves VMC, Wang Y, Schaffert RE, Hoekenga OA, Piñeros MA, Shaff JE, Klein PE, Carneiro NP, Coelho CM, Trick HN, Kochian LV (2007) A gene in the multidrug and toxic compound extrusion (MATE) family confers aluminum tolerance in sorghum. Nature genetics 39: 1156–1161

Maron LG, Piñeros MA, Guimarães CT, Magalhaes JV, Pleiman JK, Mao C, Shaff J, Belicuas SN, Kochian LV (2010) Two functionally distinct members of the MATE (multi-drug and toxic compound extrusion) family of transporters potentially underlie two major aluminum tolerance QTLs in maize. The Plant journal : for cell and molecular biology 61: 728–740

Ohyama Y, Ito H, Kobayashi Y, Ikka T, Morita A, Kobayashi M, Imaizumi R, Aoki T, Komatsu K, Sakata Y, Iuchi S, Koyama H (2013) Characterization of AtSTOP1 orthologous genes in tobacco and other plant species. Plant physiology 162: 1937–1946

Ranjan A, Sinha R, Sharma TR, Pattanayak A, Singh AK (2021) Alleviating aluminum toxicity in plants: Implications of reactive oxygen species signaling and crosstalk with other signaling pathways. Physiologia Plantarum 173: 1765–1784

Sasaki T, Ryan PR, Delhaize E, Hebb DM, Ogihara Y, Kawaura K, Noda K, Kojima T, Toyoda A, Matsumoto H, Yamamoto Y (2006) Sequence upstream of the wheat (Triticum aestivum L.) ALMT1 gene and its relationship to aluminum resistance. Plant and Cell Physiology 47: 1343–1354

Sawaki Y, Iuchi S, Kobayashi Y, Kobayashi Y, Ikka T, Sakurai N, Fujita M, Shinozaki K, Shibata D, Kobayashi M, Koyama H (2009) STOP1 regulates multiple genes that protect arabidopsis from proton and aluminum toxicities. Plant physiology 150: 281–294

Sawaki Y, Kobayashi Y, Kihara-Doi T, Nishikubo N, Kawazu T, Kobayashi M, Kobayashi Y, Iuchi S, Koyama H, Sato S (2014) Identification of a STOP1-like protein in Eucalyptus that regulates transcription of Al tolerance genes. Plant science : an international journal of experimental plant biology 223: 8–15

Silva-Navas J, Salvador N, Del Pozo JC, Benito C, Gallego FJ (2021) The rye transcription factor ScSTOP1 regulates the tolerance to aluminum by activating the ALMT1 transporter. Plant science : an international journal of experimental plant biology 310: 110951

Tsutsui T, Yamaji N, Feng Ma J (2011) Identification of a cis-acting element of ART1, a C2H2-type zinc-finger transcription factor for aluminum tolerance in rice. Plant physiology 156: 925–931

Uexküll H, Mutert E (1995) Global extent, development and economic impact of acid soils. Plant and Soil 171: 1–15

Wei Y, Han R, Xie Y, Jiang C, Yu Y (2021) Recent advances in understanding mechanisms of plant tolerance and response to aluminum toxicity. Sustainability 13: 1782

Wu L, Guo Y, Cai S, Kuang L, Shen Q, Wu D, Zhang G (2020) The zinc finger transcription factor ATF1 regulates aluminum tolerance in barley. Journal of experimental botany 71: 6512–6523

Wu W, Lin Y, Chen Q, Peng W, Peng J, Tian J, Liang C, Liao H (2018) Functional Conservation and Divergence of Soybean GmSTOP1 Members in Proton and Aluminum Tolerance. Frontiers in plant science 9: 570

Wu Y, Yang Z, How J, Xu H, Chen L, Li K (2017) Overexpression of a peroxidase gene (AtPrx64) of Arabidopsis thaliana in tobacco improves plant’s tolerance to aluminum stress. Plant molecular biology 95: 157–168

Yamaji N, Huang CF, Nagao S, Yano M, Sato Y, Nagamura Y, Ma JF (2009) A zinc finger transcription factor ART1 regulates multiple genes implicated in aluminum tolerance in rice. The Plant cell 21: 3339–3349

Ye JY, Tian WH, Zhou M, Zhu QY, Du WX, Zhu YX, Liu XX, Lin XY, Zheng SJ, Jin CW (2021) STOP1 activates NRT1.1-mediated nitrate uptake to create a favorable rhizospheric pH for plant adaptation to acidity. Plant Cell 12: 3658–3674

Yokosho K, Yamaji N, Fujii-Kashino M, Ma JF (2016) Retrotransposon-Mediated Aluminum Tolerance through Enhanced Expression of the Citrate Transporter OsFRDL4. Plant Physiology 172: 2327–2336

Zhang Y, Zhang J, Guo J, Zhou F, Singh S, Xu X, Xie Q, Yang Z, Huang C (2019) F-box protein RAE1 regulates the stability of the aluminum-resistance transcription factor STOP1 in Arabidopsis. Proceedings of the National Academy of Sciences 116: 319–327

Zhou F, Singh S, Zhang J, Fang Q, Li C, Wang J, Zhao C, Wang P, Huang CF (2023) The MEKK1-MKK1/2-MPK4 cascade phosphorylates and stabilizes STOP1 to confer aluminum resistance in Arabidopsis. Molecular plant 16: 337–353

